# Plastic Leachate Exposure Drives Antibiotic Resistance and Virulence in Marine Bacterial Communities

**DOI:** 10.1101/2023.02.13.528379

**Authors:** Eric J. Vlaanderen, Timothy M. Ghaly, Lisa R. Moore, Amaranta Focardi, Ian T. Paulsen, Sasha G. Tetu

## Abstract

Plastic pollution is a serious global problem, with more than 12 million tonnes of plastic waste entering the oceans every year. Plastic debris can have considerable impacts on microbial community structure and functions in marine environments, and has been associated with an enrichment in pathogenic bacteria and antimicrobial resistance (AMR) genes. However, our understanding of these impacts is largely restricted to microbial assemblages on plastic surfaces. It is therefore unclear whether these effects are driven by the surface properties of plastics, providing an additional niche for certain microbes residing in biofilms, and/or chemicals leached from plastics, the effects of which could extend to surrounding planktonic bacteria. Here, we examine the effects of polyvinyl chloride (PVC) plastic leachate exposure on the relative abundance of genes associated with bacterial pathogenicity and AMR within a seawater microcosm community. We show that PVC leachate, in the absence of plastic surfaces, drives an enrichment in AMR and virulence genes. In particular, leachate exposure significantly enriches AMR genes that confer multidrug, aminoglycoside and peptide antibiotic resistance. Additionally, enrichment of genes involved in the extracellular secretion of virulence proteins was observed among pathogens of marine organisms. This study provides the first evidence that chemicals leached from plastic particles alone can enrich genes related to microbial pathogenesis within a bacterial community, expanding our knowledge of the environmental impacts of plastic pollution with potential consequences for human and ecosystem health.

## 1. Introduction

Plastic pollution in marine ecosystems has become a serious global problem, with more than 12 million metric tonnes of plastic waste ending up in the oceans every year (Borrelle et al., 2020; Jambeck et al., 2015; Lau et al., 2020). As plastic production rates continue to rise and poor waste management practices remain in many areas of the world, issues associated with marine plastic pollution are likely to increase in the future (Borrelle et al., 2020; Jadhav et al., 2022; Lau et al., 2020; Lebreton and Andrady, 2019). To date, recognition of the environmental impacts of plastic debris has largely focused on entanglement and ingestion by marine species (Cózar et al., 2014; Gregory, 2009; Lebreton et al., 2018; Wright et al., 2013). However, additional impacts are increasingly being recognised for marine microorganisms.

These include the environmental release of chemicals via leaching from plastic particles, which can significantly alter marine microbial communities (Capolupo et al., 2020; Focardi et al., 2022; Gunaalan et al., 2020; Romera-Castillo et al., 2018) and dissemination of pathogenic microorganisms, via rafting on plastic debris (Bhagwat et al., 2021; Bryant et al., 2016; Zettler et al., 2013; Zhang et al., 2022).

Plastics are known to leach a variety of organic and inorganic substances through weathering and biological degradation processes. This includes additives such as plasticizers, UV stabilizers, metals, and dyes, most of which are not chemically bound to the polymer matrix (Hahladakis et al., 2018; Hermabessiere et al., 2017). Some of these additives are known endocrine-disrupters, reproductive toxicants, carcinogens, and mutagens (Wiesinger et al., 2021; Zimmermann et al., 2019). The ability to tolerate exposure to common inorganic and/or organic components of plastic leachate has been shown to be highly variable across different marine microbes. Exposure to chemicals leaching from plastics is detrimental to some marine organisms, such as zooplankton (Gewert et al., 2021; Lithner et al., 2009), green algae (Simon et al., 2021), bacterial picocyanobacteria (Sarker et al., 2020; Tetu et al., 2019), and other keystone marine microbes, including SAR11 (Focardi et al., 2022). However, some marine heterotrophic bacteria appear to benefit from plastic leach exposure, likely from the increase in available dissolved organic carbon (Birnstiel et al., 2022; Focardi et al., 2022; Romera-Castillo et al., 2018).

Colonisation of plastic debris by microorganisms, termed the “plastisphere”, has been extensively studied, with clear indications that this niche selects for microbial communities differing in abundance and diversity from the surrounding waters (Bryant et al., 2016; Dussud et al., 2018; He et al., 2022; Zettler et al., 2013). Of particular concern is the enrichment of potential pathogens and antibiotic resistant microbes on plastic particles, as well as increases in antimicrobial resistance (AMR) genes (Di Pippo et al., 2022; Loiseau and Sorci, 2022; Oberbeckmann et al., 2015; Sathicq et al., 2021; Sucato et al., 2021; Sun et al., 2021; Wang et al., 2020; Yang et al., 2019; Zhang et al., 2022). Studies using metagenomics-derived data have found higher relative abundance, diversity and richness indices of human and fish pathogens in the microbial communities attached to plastics in the Mediterranean Sea (Dussud et al., 2018), the North Pacific Gyre (Yang et al., 2019), and in coastal regions of Norway (Radisic et al., 2020), the Gulf of Mexico (Sun et al., 2021), and Eastern Australia (Bhagwat et al., 2021).

It is clear that marine plastic debris has the potential to serve as a vector for pathogens and genes involved in virulence and antibiotic resistance. However, it is not clear whether this is due solely to plastic debris providing an additional niche for certain microbes, particularly those residing in biofilms, or because leached plastic chemicals also favour increases in such microorganisms. To our knowledge there have been no studies examining whether chemicals that leach from plastic waste select for higher relative abundances of AMR and pathogenicity traits within a community, independent of the physical effects linked to plastic particles. Here we demonstrate that polyvinyl chloride (PVC) plastic leachate enriches for virulence and AMR genes in a marine microbial community from Eastern Australian coastal shelf waters (Focardi et al., 2022).

## 2. Methods

### 2.1. Data acquisition

Metagenomic data was obtained from our previous study examining the effects of PVC leachate and zinc, an abundant PVC additive, on a seawater microcosm community (Focardi et al., 2022). Methodology for the leachate preparation, and microcosm experiment set-up has been described in our previous study (Focardi et al., 2022). Briefly, microcosm samples were subject to a six-day exposure to either 1% (0.5g/L) PVC leachate (PVC1), 10% (5g/L) PVC leachate (PVC10), 0.13 mg/L zinc chloride (ZnL), 1.3 mg/mL zinc chloride (ZnH) (zinc being the most abundant inorganic component in leachate from this PVC plastic), or left untreated as control samples (SW). DNA extracted from each treatment was used to generate metagenomic libraries at the Ramaciotti Center for Genomics (Sydney, Australia) using the Illumina Nextera DNA Flex library preparation kit, and sequenced on the NovaSeq6000 platform (2×150 bp High Output run). Gene sequences, de-replicated at 98% nucleotide identity, and gene counts for each sample were retrieved from Focardi et al. (2022). Detailed methodology for the metagenomic data processing, assembly, gene prediction, and gene counts has been described in our previous study. All raw sequence data are available under NCBI BioProject accession PRJNA756323.

### 2.2. Identification and quantification of AMR, virulence, and toxin genes

AMR and pathogenicity-related genes from each sample were identified using PathoFact v1.0 (de Nies et al., 2021) with default settings. PathoFact is a pipeline that identifies putative virulence factors, bacterial toxins, and AMR genes. PathoFact further classifies AMR genes by antimicrobial category and resistance mechanism. For cases where AMR genes were assigned multiple resistance mechanisms, we used only the first predicted mechanism from PathoFact.

Relative abundance of each gene was estimated with normalised read counts using the simplified transcripts per million (TPM) method described by Wagner et al. (2012). The per-sample mean number of nucleotides mapped per feature was taken as a proxy for read length. One-way ANOVAs followed by post-hoc Tukey-HSD tests were performed to compare the relative abundance of genes in PVC treatment samples (PVC1 and PVC10) and Zn treatment samples (ZnL and ZnH) independently against the control samples (SW) in each of the PathoFact output categories (AMR, Virulence, and Toxin).

### 2.3. Additional screening and categorisation of virulence genes

PVC leachate treatment resulted in increases in the relative abundance of virulence genes that were deemed significant, but were close to the significance cut-off (P-value < 0.05) as predicted by PathoFact. Thus, to investigate this further, we used a second analysis pipeline SeqScreen v3.4 (Balaji et al., 2022) with the SeqScreenDB v2.0 (version February 2022) to test whether increases in the relative abundance of virulence genes were significant.

SeqScreen utilises a large set of curated Functions of Sequences of Concern (FunSoCs) specific to microbial pathogenesis to both identify and characterise virulence genes. We employed SeqScreen [parameters: --splitby 50000 --threads 24] to re-screen all metagenomic data for virulence genes and determine which virulence functional categories these genes were assigned to. Since our goal in using SeqScreen was specifically to identify bacterial virulence factors, genes assigned to virus-specific categories or AMR were removed from the SeqScreen output. One-way ANOVAs followed by post-hoc Tukey-HSD tests were performed to compare the relative abundance of virulence genes among all treatment and control groups.

### 2.4. Enrichment of specific AMR/virulence categories and genes

For PathoFact-predicted AMR genes, we compared the difference in relative abundance of antimicrobial categories and resistance mechanisms between PVC treatments and control samples. For SeqScreen-predicted virulence genes, we compared the difference in relative abundance of bacterial virulence FunSoC categories. Categories which had a mean relative abundance across PVC and seawater samples below 100 TPM for virulence and 10 TPM for AMR were excluded from the results as we considered them unlikely to be biologically relevant. For all comparisons, normalised gene counts were summed by category for each sample. One-way ANOVA tests were run for each category and followed by post-hoc Tukey-HSD tests where significant results were identified.

The set of predicted AMR genes (PathoFact) and virulence genes (SeqScreen) most highly enriched among PVC leachate-treated communities were then identified for further analysis. The genes most enriched in the PVC10 treatment compared to the seawater control were identified by calculating the log2-fold change of genes with a minimum mean relative abundance of 9 TPM for AMR, and 20 TPM for virulence in the treatment group (in order to select genes that were both highly abundant and highly enriched). We assigned putative taxonomy to the 20 most highly enriched AMR genes using a BLASTn search against the

NCBI nt database. For virulence genes, we used the taxonomic assignments provided by SeqScreen, which runs both a DIAMOND (Buchfink et al., 2015) search against a curated UniRef100 database (Suzek et al., 2007), as well as running Centrifuge (Kim et al., 2016) against Archaeal and Bacterial RefSeq genomes.

### 2.5. Diversity of AMR and virulence genes

Beta-diversity of AMR (PathoFact) and virulence (SeqScreen) gene profiles between treated and untreated microbial communities were visualised using a non-metric muti-dimensional scaling (nMDS) plot based on the Bray-Curtis index of normalised read counts. This was achieved using the *vegdist* and *metaMDS* functions of the VEGAN v2.5-7 package (Oksanen et al., 2013) in R (v 4.1.2). Significant differences between treatment groups were analysed using multivariate PERMANOVA using the *adonis2* function in VEGAN.

Alpha diversity was calculated in R using Shannon-Weiner and Simpson indexes for the AMR (PathoFact) and virulence (SeqScreen) gene sets. One-way ANOVA and post-hoc Tukey-HSD tests were run against the resulting values across PVC1, PVC10 and SW.

## 3. Results and Discussion

Previously, we examined the effects of exposure to two concentrations of PVC plastic leachate and of zinc, the most abundant inorganic PVC additive, on a marine microbial community via a six-day microcosm experiment, showing that this leads to substantial changes in community composition and function (Focardi et al., 2022). In this study, we have analysed the metagenomic data from Focardi et al. (2022), to investigate whether exposure to PVC leachate and/or zinc results in significant enrichment of genes associated with pathogenicity and drug resistance.

### 3.1. Plastic leachate exposure increases antibiotic resistance and virulence genes in marine microbial communities

Marine microbial communities treated with PVC leachate, in the absence of physical plastic surfaces, showed a concentration-dependent increase in the relative abundance of AMR, virulence and toxin genes compared to non-treated controls, although the increase in toxin genes was not statistically significant (ANOVA, p=0.082, Suppl. Table 1f) (Fig 1a-c, based on PathoFact predictions). Past analyses of leachate from this specific PVC plastic showed it is comprised of a complex mix of both organic and inorganic substances, with levels of zinc, a common PVC additive, found to be particularly high (Tetu et al., 2019). As Zn exposure has previously been shown to increase the prevalence of antibiotic resistant bacteria in the environment (e.g., Poole, 2017; Silva et al., 2021), and promote virulence in host-associated bacteria (Wu et al., 2021), we also looked to see if exposure to zinc alone was sufficient to account for the PVC leachate impact on AMR and virulence gene prevalence. Using PathoFact predictions, we found that treatment with two concentrations of zinc, ZnL (0.13 mg/L ZnCl) and ZnH (1.3 mg/mL ZnC1), had no effect on the relative abundance of AMR, virulence or toxicity genes compared to untreated seawater controls (Fig. 1d-f). This suggests that zinc additives alone are not driving the observed effects of PVC leachate on AMR and virulence gene enrichment.

**Fig. 1.**
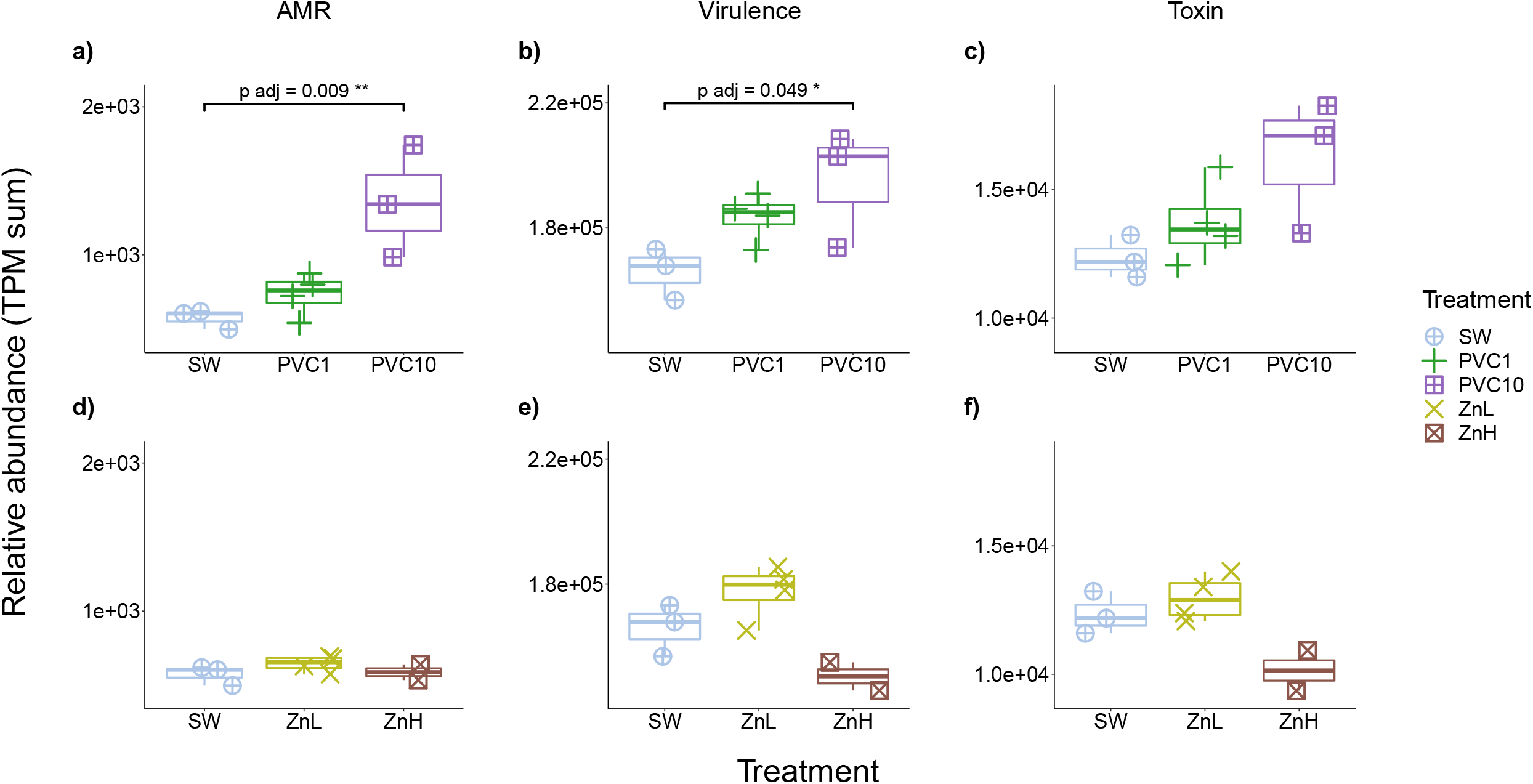
Relative abundance (TPM sum) of genes encoding; (a, d) AMR, (b, e) virulence, and (c, f) toxin functions, predicted by PathoFact for 1% PVC leachate (PVC1), 10% PVC leachate (PVC10), 0.13 mg/L zinc chloride (ZnL), and 1.3 mg/mL zinc chloride (ZnH) treatments, compared with seawater (SW). Tukey-HSD adjusted p-values have been reported for treatments which differ significantly from the control. The full set of statistical results for these tests are provided in Supplementary Table 1(b-m).

AMR gene relative abundance showed a slight, non-significant increase in the 1% PVC leachate treatments and a strong, significant increase in the 10% PVC leachate treatment relative to untreated seawater controls (Fig. 1a; 2.4-fold increase; TukeyHSD, p-adj.=0.009, Suppl. Table 1a, 1c). Similarly, for the set of virulence-associated genes based on PathoFact predictions, 1% PVC leachate treatment resulted in a small non-significant increased while the 10% PVC leachate treatment drove a significant increase in virulence genes (Fig. 1b; 1.2-fold increase; TukeyHSD, p-adj.=0.049, Suppl. Table 1a, 1e). Given that this increase was close to the significance cut-off, we performed further analysis of virulence genes, using the recently developed SeqScreen pipeline which has a larger, curated virulence database (Balaji et al., 2022). Based on this, both 1% and 10% PVC leachate treatments resulted in significant enrichments of virulence genes (Suppl. Fig. 1, Tukey-HSD, p adj. = 0.022 and p adj. = 0.004, respectively, Suppl. Table 5b), representing a 1.3-fold increase in virulence genes in the PVC1 and 1.4-fold for PVC10 (Suppl. Table 5a).

### 3.2. Plastic leachate exposure changes the makeup of AMR gene suites and resistance mechanisms

Both 1% and 10% PVC leachate treatments drove clear shifts in AMR gene profiles, evident from non-metric multidimensional scaling (NMDS) analysis (Fig. 2a, stress value = 0.06, indicating clear separation from both the control and zinc treatments) and supported by PERMANOVA (p=0.001, R^2^ = 0.57, Suppl. Table 2a). However, PVC treatments had no significant effect on the alpha diversity of AMR genes (Fig. 2b, c), indicating that enrichment of AMR genes following PVC leachate exposure is due to an increase in the relative abundance of specific AMR genes, rather than an increase in overall AMR gene diversity.

**Fig. 2.**
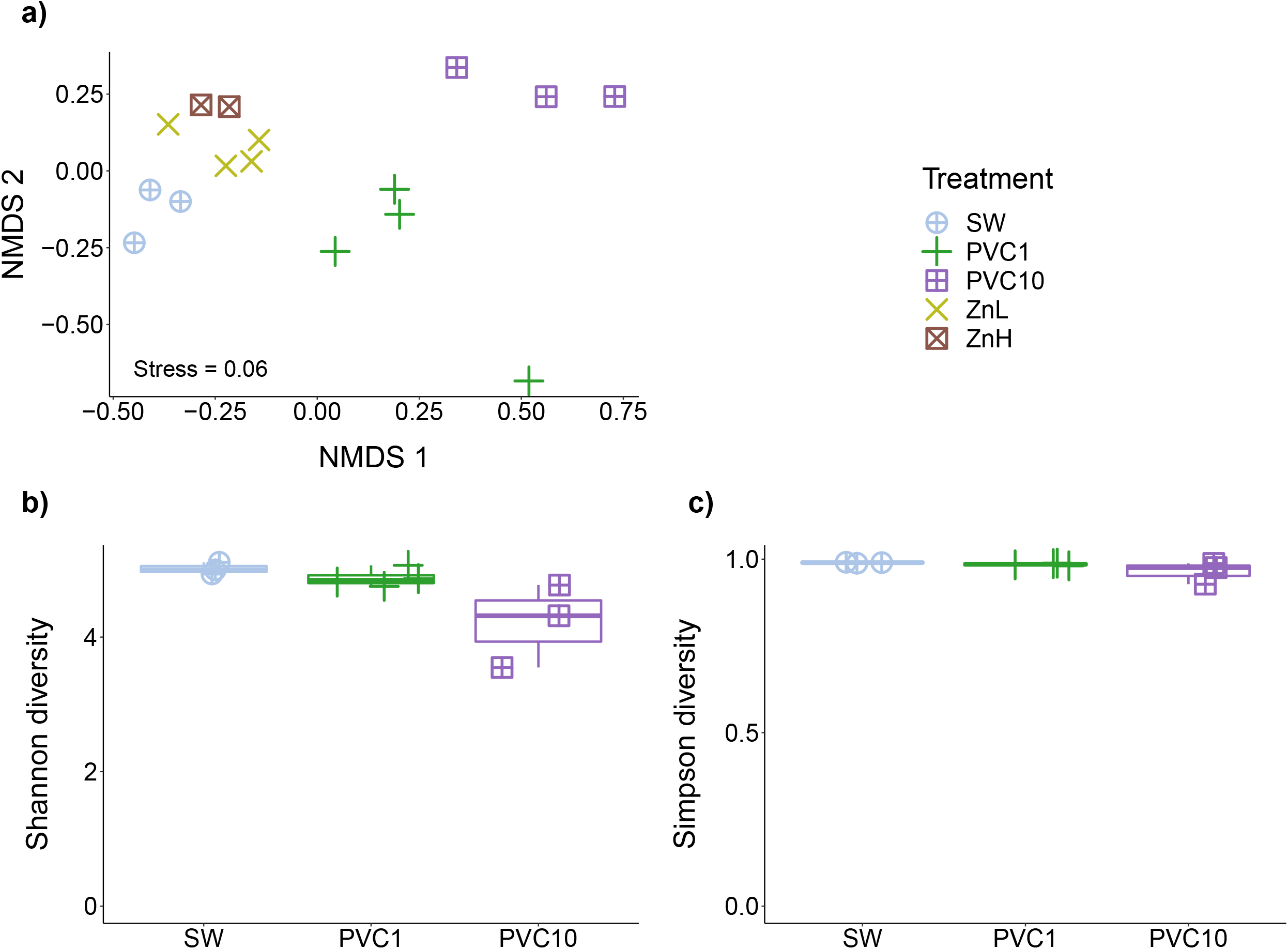
a) NMDS plot of AMR gene profiles for all samples, b) Shannon-Wiener, and c) Simpson diversity of AMR genes for seawater controls (SW), and 1% PVC leachate (PVC1), and 10% PVC leachate (PVC10) treatments.

The 10% PVC leachate treatment drove significant enrichments in several AMR categories (Fig. 3a), including aminoglycoside (Tukey-HSD, p adj _≤_ 0.0001), antimicrobial peptide (Tukey-HSD, p adj=0.001), aminoglycoside:aminocoumarin (Tukey-HSD, p adj=0.02), and multidrug resistance (Tukey-HSD, p adj=0.02). Aminoglycoside resistance genes were also significantly enriched following the 1% PVC leachate treatment (Tukey-HSD, p adj=0.02). In contrast, genes belonging to the MLS category (macrolides, lincosamides, and streptogramins) were significantly lower in abundance in both 1% and 10% PVC leachate treatments compared to the seawater control (Tukey-HSD, p adj=0.04 and p adj=0.005 respectively) (Suppl. Table 3b). As MLS antibiotics are primarily active against Gram positive bacteria, these resistance genes are typically found in these organisms. Thus, this decline in MLS resistance gene abundance may be due to the large increase in relative abundance of Gram negative bacteria following leachate exposure (Focardi et al., 2022).

**Fig. 3.**
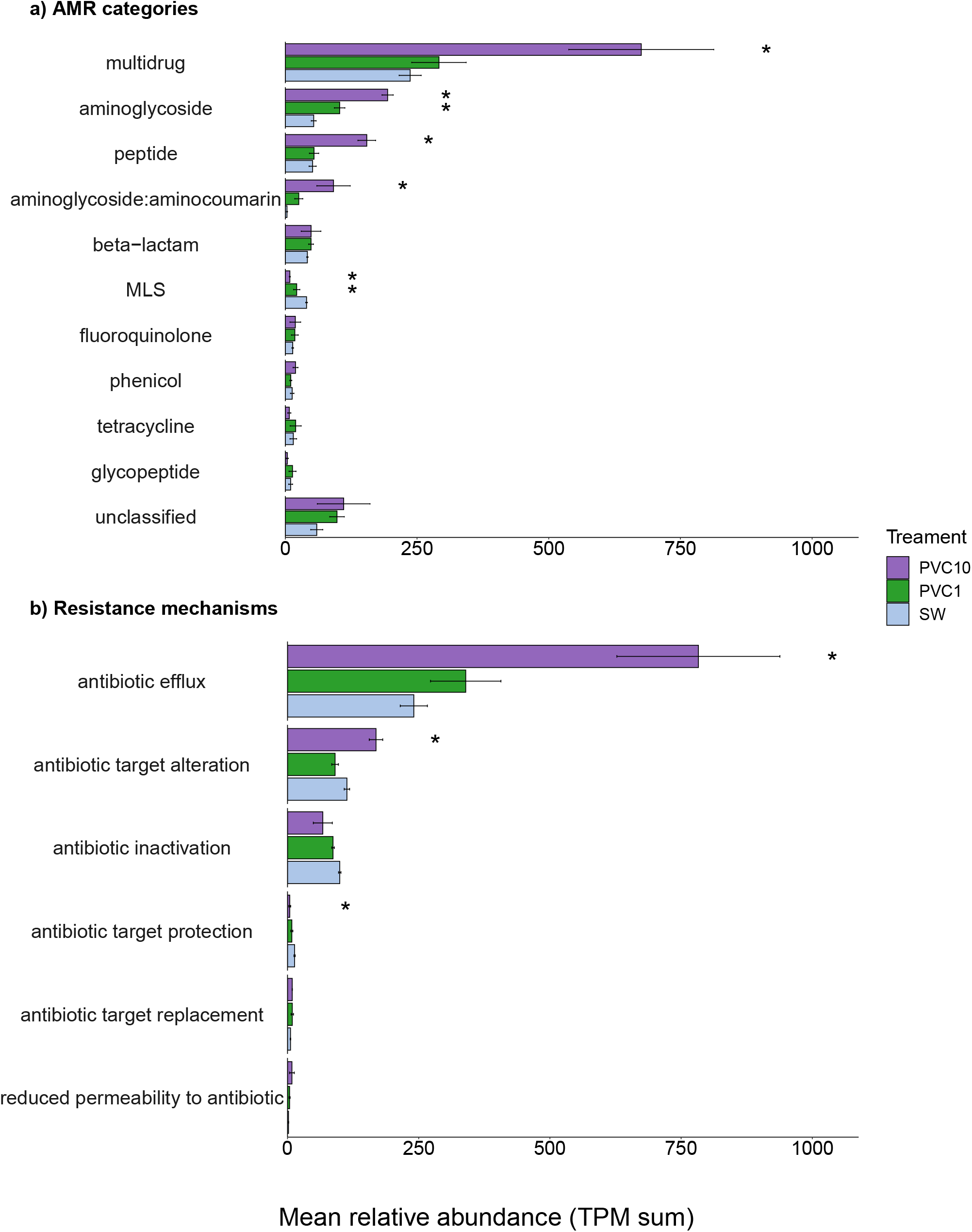
Comparison of the mean relative abundance (TPM sum) between PVC and seawater samples for a) antimicrobial resistance categories and b) antibiotic resistance mechanisms. Error bars indicate the standard error of the mean and stars (*) denote a significant difference from the control (SW) (Tukey-HSD, p < 0.05). Full statistical results for tests displayed here are provided in Supplementary Tables 3 and 4.

The profiles of resistance mechanisms within PVC leachate-treated communities were also altered (Fig. 3b). In particular, the 10% PVC leachate treatment led to a significant enrichment of genes that confer AMR via antibiotic efflux (Tukey-HSD, p=0.01) and antibiotic target alteration (Tukey-HSD, p<0.01) mechanisms compared to untreated seawater (Suppl. Table 4b). These two resistance mechanism categories encompass the majority of AMR genes identified in this study (46% assigned to antibiotic efflux, 21% assigned to antibiotic target alteration). Resistance genes assigned to the antibiotic target protection category were significantly lower in abundance in 10% PVC compared to seawater, however, this is a small category with fewer than 4% of AMR genes assigned to it overall.

The twenty most enriched AMR genes, showing the highest fold change in the 10% PVC leachate treatment, were examined to determine their likely host organism and which AMR category and resistance mechanism each was assigned to (Table 1).

**Table 1.** Characteristics of the most highly enriched AMR genes following 10% PVC exposure.^1^

| AMR Gene | Gene ID | Fold Change<br>PVC10:SW<br>(log<sub>2</sub>) | Mean<br>Relative<br>Abundance<br>in PVC10<br>(TPM) | AMR<br>Category | Resistance<br>Mechanism | Predicted genus |
| --- | --- | --- | --- | --- | --- | --- |
| *ampC* | c_000000000144_14 | ∞ | 10.8 ± 7 | beta-lactam | Antibiotic inactivation | *Tritonibacter* |
| *emrE* | c_000000000007_139 | ∞ | 9.6 ± 6.5 | multidrug | Antibiotic efflux | *Tritonibacter* |
| *mtrE* | c_000000000080_44 | ∞ | 9.4 ± 5.3 | multidrug | Antibiotic efflux | *Tritonibacter* |
| *ugd* | c_000000000038_92 | 8.8 | 14.4 ± 9.8 | peptide | Antibiotic target alteration | *Tritonibacter* |
| *abeS* | c_000000000316_5 | 7.6 | 22.7 ± 17.4 | multidrug | Antibiotic efflux | *Pseudoalteromonas* |
| *acrB* | c_000000000018_83 | 6.8 | 77.6 ± 45.9 | multidrug | Antibiotic efflux | *Alteromonas* |
| *ugd* | c_000000000027_138 | 6.8 | 64.6 ± 39.3 | peptide | Antibiotic target alteration | *Alteromonas* |
| *ksgA* | c_000000000010_130 | 6.8 | 69.3 ± 41.1 | aminoglycoside | - | *Alteromonas* |
| *mexT* | c_000000000025_20 | 6.7 | 56.7 ± 34.5 | multidrug | Antibiotic efflux | *Alteromonas* |
| *adeF* | c_000000000005_64 | 6.6 | 66.9 ± 39 | multidrug | Antibiotic efflux | *Alteromonas* |
| *baeR* | c_000000000073_7 | 6 | 60.8 ± 37.1 | aminoglycoside:<br>aminocoumarin | Antibiotic efflux | *Alteromonas* |
| *mexT* | c_000000001693_3 | 5.9 | 17.9 ± 14.5 | multidrug | Antibiotic efflux | *Alteromonas* |
| *ksgA* | c_000000011168_2 | 5.3 | 16.3 ± 12.8 | aminoglycoside | - | *Paraglaciecola* |
| *mexT* | c_000000009727_1 | 5.2 | 19.8 ± 13.8 | multidrug | Antibiotic efflux | *Alteromonas* |
| *pmpM* | c_000000003002_4 | 4.1 | 10.7 ± 6.7 | multidrug | Antibiotic efflux | *Alcanivorax* |
| *adeF* | c_000000085939_101 | 4 | 10.5 ± 6.4 | multidrug | Antibiotic efflux | *Alcanivorax* |
| *qepA* | c_000000000289_9 | 3.8 | 13 ± 8 | fluoroquinolone | - | *Alcanivorax* |
| *ugd* | c_000000006705_2 | 3.7 | 20.5 ± 15 | peptide | Antibiotic target<br>alteration | *Vibrio* |
| *crp* | c_000000000481_8 | 3.7 | 11.3 ± 6.9 | unclassified | - | *Alcanivorax* |
| *baeR* | c_000000005224_3 | 3.7 | 13.4 ± 8.2 | aminoglycoside:<br>aminocoumarin | Antibiotic efflux | *Alcanivorax* |

<sup>†</sup>The AMR gene, category, resistance mechanism (as provided by PathoFact), and predicted genus (based on NCBI BLASTn) for AMR genes which were found to be highly enriched in 10% PVC treatments, sorted by fold change (log<sub>2</sub>). Fold change has been reported as ∞ for genes which were not observed in the seawater community.

Twelve out of the twenty most enriched antibiotic resistance genes are related to antibiotic efflux. These include efflux pumps from the RND, SMR and MATE multidrug efflux pump families, as well as MexT and BaeR, which are regulators of RND efflux pump gene expression (Henderson et al., 2021). All three of these efflux pump families typically have broad substrate specificities, particularly the RND efflux pumps. In addition to antibiotics, these efflux pumps can often export a wide range of complex hydrophobic organic molecules. Thus, it is possible that the increased abundance of efflux pumps following exposure to plastic leachate may be due to their ability to protect against toxic organic components in the leachate, exporting such components out of the cell. Of the remaining most abundant resistance genes, three are target site alteration and all of these are *ugd* genes, which provide polymyxin resistance via lipopolysaccharide modification, and the beta-lactamase gene, *ampC*, involved in antibiotic inactivation.

The most highly enriched AMR genes are all predicted to be found in heterotrophic, predominantly Gram negative bacteria in the microcosms, with *Tritonibacter, Alteromonas* and *Alcanivorax* the most common predicted hosts of these genes. This is consistent with what taxonomic groups were observed to be most enriched in the PVC treated samples (Focardi et al., 2022). *Tritonibacter* are marine bacteria, originally described from a cultured representative isolated from oil-contaminated surface water during the Deepwater Horizon oil spill (Klotz et al., 2018). *Alteromonas* has been reported to be one of the main groups of microbes capable of growing in plastic leachates (Birnstiel et al., 2022). *Alcanivorax* are alkane degrading marine bacteria that are found in low abundance in surface marine waters but are highly enriched in oil contaminated marine environments (Hara et al., 2003) and have previously been reported to encode multiple multidrug resistance proteins (Sinha et al., 2021).

While none of the genera containing these abundant AMR genes include known human pathogens, with the exception of *Vibrio*, there is potential for gene transfer events, facilitated by mobile genetic elements, to move AMR genes between lineages, and into species which may pose a risk to human health. At least in *Escherichia coli*, plastic leachate has been shown to upregulate horizontal gene transfer (Yuan et al., 2022), opening the possibility of synergistic effects that enrich bacteria harbouring AMR genes, whilst also facilitating AMR spread. Indeed, capture of AMR genes by mobile genetic elements has been well documented, and in many cases, has resulted in their spread into diverse human pathogens across the globe, originating from single mobilisation events (Moellering Jr, 2010; Wang et al., 2018). Further, a large proportion of AMR genes now globally circulating among clinical pathogens are predicted to have originated in marine environments, including several efflux pump and beta-lactamase genes (Ghaly et al., 2021). *Alteromonas* species harbour large conjugative elements that can facilitate this movement, such as mega-plasmids and integrative and conjugative elements (ICEs) (Cusick et al., 2020; López-Pérez et al., 2017). In fact, several characterised ICEs are shared between *Alteromonas* and human pathogens (López-Pérez et al., 2017; Pang et al., 2016), indicating the potential transmission of genes from environmental to clinical organisms.

### 3.3. Plastic leachate exposure changes the composition of virulence factors

PVC leachate treatments drove clear shifts in virulence gene profiles (SeqScreen-derived), evident from NMDS analysis (Fig. 4a, stress value = 0.05, indicating clear separation from both the control and zinc treatments) and supported by PERMANOVA (p=0.001, R^2^ = 0.59, Suppl. Table 6a). PVC 10% treatment had a significant negative effect on the Shannon-Wiener diversity of virulence genes (Fig. 4b; Tukey-HSD, p adj = 0.004, Suppl. Table 6c), however, no such effect was observed on Simpson diversity (Fig 4c). This indicates that enrichment of virulence genes following PVC leachate exposure is due to an increase in the relative abundance of specific virulence genes, rather than an increase in overall virulence gene diversity.

**Fig. 4.**
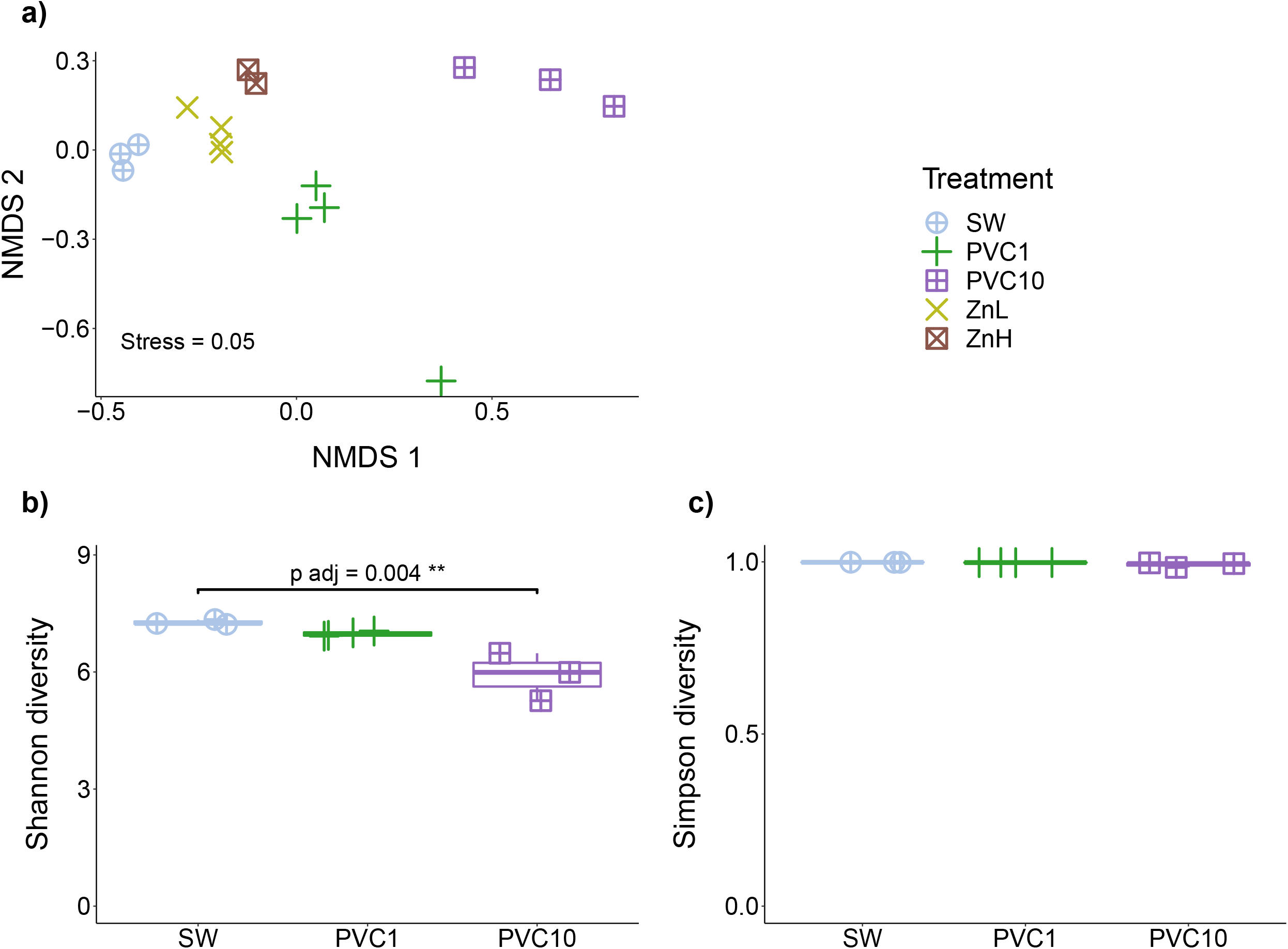
a) NMDS plot of virulence gene profiles from SeqScreen for all samples, b) Shannon-Wiener, and c) Simpson diversity of virulence genes for seawater controls (SW), and 1% PVC leachate (PVC1), and 10% PVC leachate (PVC10) treatments.

PVC leachate treatments led to changes in the composition of virulence genes, based on SeqScreen assigned virulence categories (Fig. 5). Both 1% and 10% PVC treatments resulted in significant increases in the relative abundance of two virulence categories: secretion (Tukey-HSD, p-adj = 0.003 for PVC1, p-adj < 0.0001 for PVC10), and bacterial counter signalling (Tukey-HSD, p-adj = 0.026 for PVC1, p-adj < 0.0001 for PVC10) (Suppl. Table 7b). The secretion category includes the components of bacterial secretion systems, and the bacterial counter signalling category includes genes involved in the suppression of host immune signalling to avoid inflammatory responses. The toxin synthase category, however, was significantly reduced in PVC10 samples (Tukey-HSD, p adj = 0.005, Suppl. Table 7b).

**Fig. 5.**
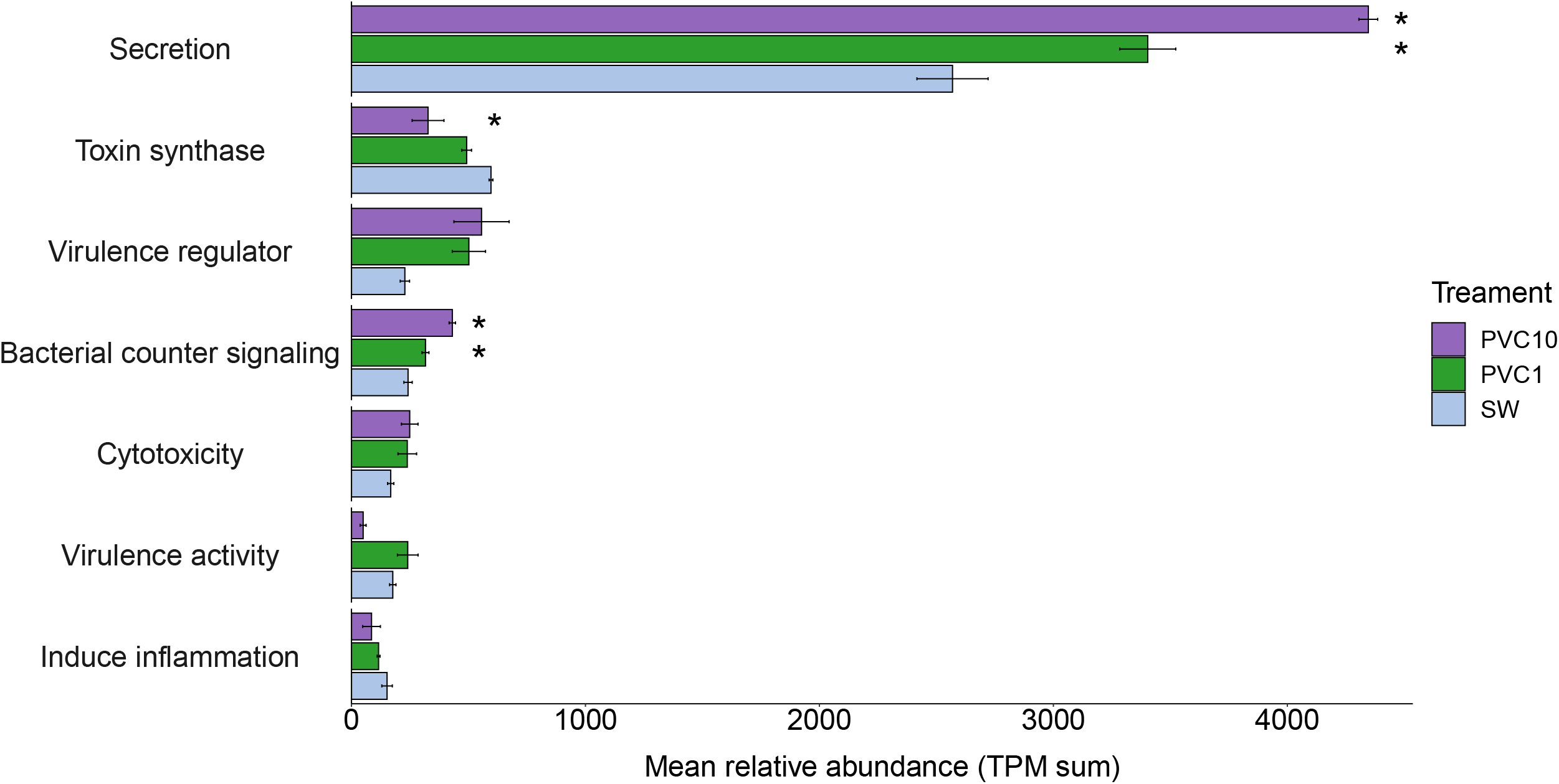
Comparison of the mean relative abundance (TPM sum) between PVC and seawater samples for virulence categories (provided by SeqScreen). Error bars indicate the standard error of the mean and stars (*) denote a significant difference from the control (SW) (Tukey-HSD, p adj < 0.05). Full statistical results for tests displayed here are provided in Supplementary Table 7.

This category includes enzymes involved in the production or modification of toxins. In the SeqScreen database, this category is largely focused on mycotoxins (those synthesised by fungi). Plastic leachate has toxic effects on fungi, impairing fungal enzymatic activity (Li et al., 2022). Thus, a decline in this category may be due to the negative effects of PVC leachate exposure on marine fungi within the seawater microcosm.

Analysis of the virulence genes most strongly enriched in the 10% PVC leachate treatment was carried out to determine their likely host organism and which virulence category each falls under. Table 2 lists the twenty genes with the highest fold change enrichment.

**Table 2.**
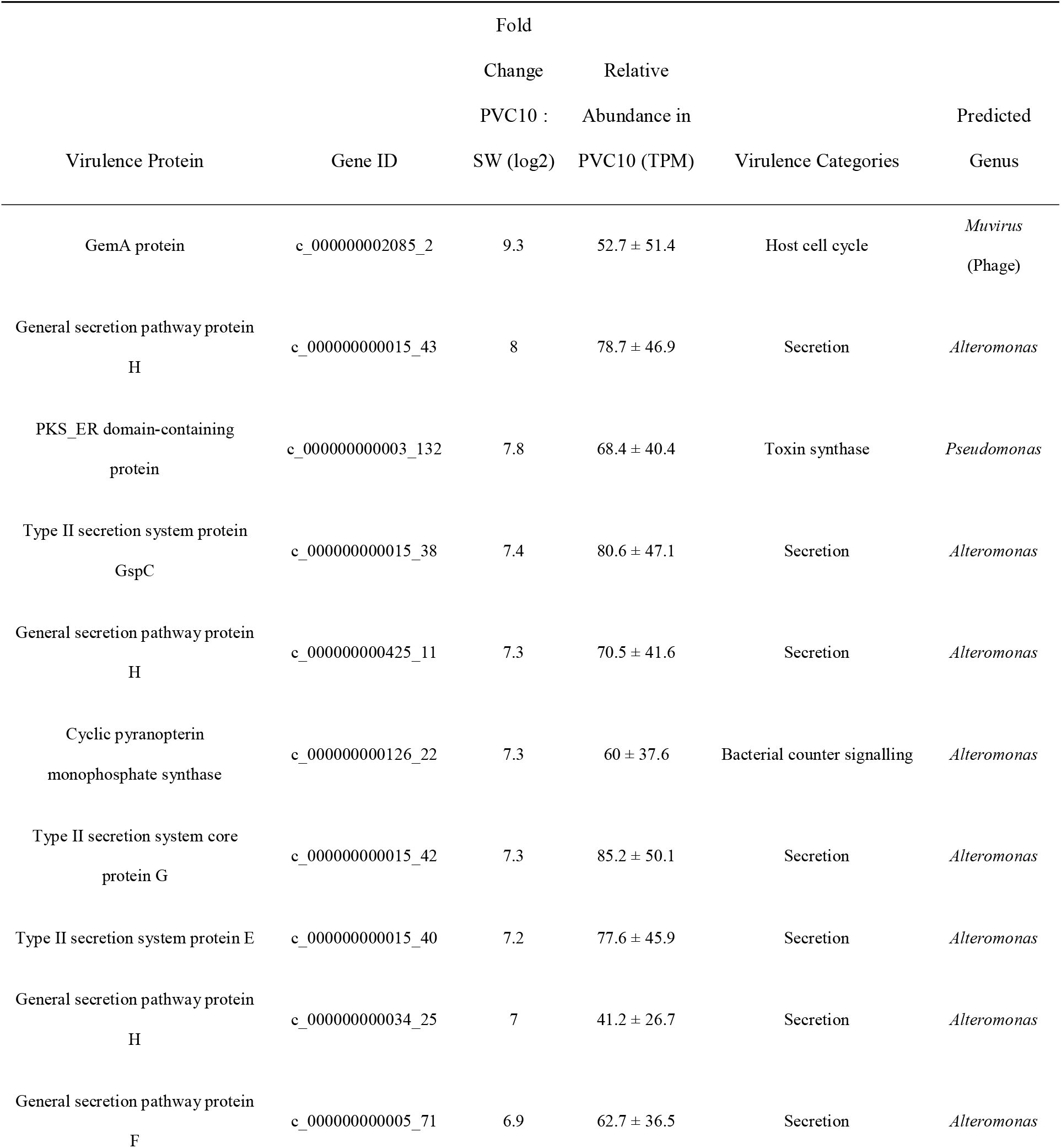
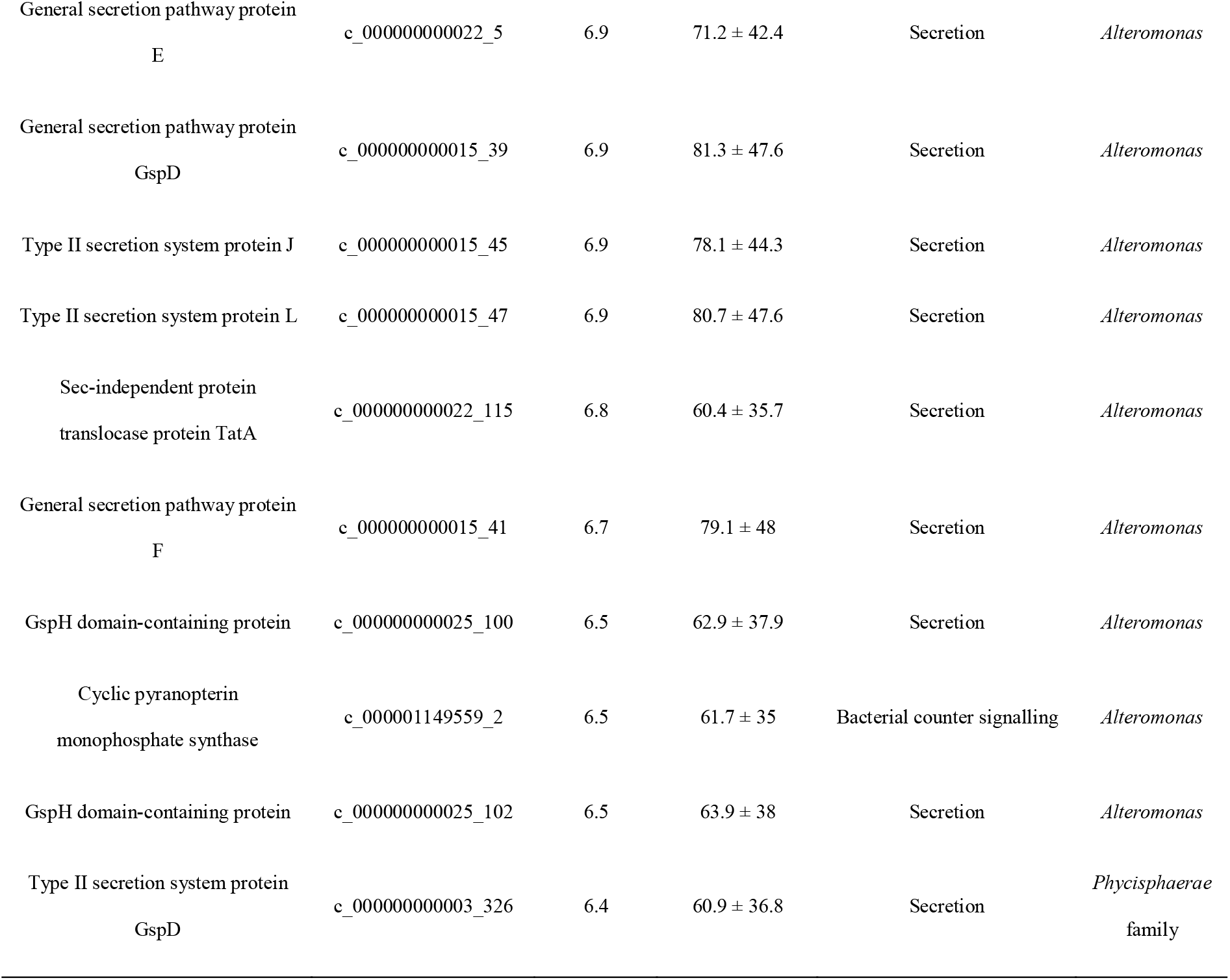
Top 20 Virulence genes enriched in 10% PVC treatments, sorted by fold change (log_2_) based on SeqScreen analyses.

Sixteen out of the twenty most enriched genes are involved in secretion, encoding components of the general secretion (Sec) pathway and Type II secretion systems (T2SSs). The Sec pathway is used by several bacterial pathogens to secrete proteins that promote their virulence (Green and Mecsas, 2016). Although the Sec pathway does not export proteins outside of the cell, in Gram negative bacteria, proteins delivered by the Sec pathway to the periplasm can be exported with the aid of T2SSs (Green and Mecsas, 2016). T2SS channels are located only in the outer membrane, and thus can only export proteins that have been delivered to the periplasm by other pathways, including the Sec pathway (Korotkov et al., 2012). Thus, the simultaneous enrichment of both T2SS and Sec pathway components suggests that PVC leachate exposure leads to an increase in microbes that employ extracellular protein secretion. Several bacterial pathogens use T2SSs to secrete proteins associated with host disease, such as hemolysins, lipases, proteases, esterases, polygalacturonases, deubiquitinases, aerolysins, DNases, amylases, and mucin-degrading enzymes (Cianciotto and White, 2017).

*Alteromonas* spp. appear to be largely responsible for driving the increase in virulence genes following PVC leachate exposure (Table 2). Several *Alteromonas* spp. have been reported as coral and algal pathogens (Brown et al., 2013; Peng and Li, 2013; Vairappan et al., 2001), and associated with disease in marine arthropods (Alfiansah et al., 2020). Plastic pollution has been shown to have toxic effects on both algae and marine invertebrates (Haegerbaeumer et al., 2019; Pisani et al., 2022; Simon et al., 2021; Zhu et al., 2022), and entanglement by plastic particles may significantly increase the risk of disease in scleractinian corals (Lamb et al., 2018). Here, we show that an additional consequence of plastic pollution for these organisms might be greater disease susceptibility due to the enrichment of pathogenic bacteria and their associated virulence traits.

## 4. Conclusion

There is growing evidence that plastic pollution in marine environments can lead to an enrichment in pathogenic bacteria and AMR genes. However, it is unclear whether these effects are driven by the physical or chemical attributes of plastic marine pollution, as differential colonisation and growth rates on plastic particles may be driven by physical surface properties and/or chemicals leached from the plastic. Here we show that PVC leachate, in the absence of plastic surfaces, drives an enrichment in AMR and virulence genes within a seawater community. The enrichment of pathogenic bacteria and virulence traits may have serious consequences for environments which are frequently exposed to human pollution, such as urban harbours and aquacultural settings. Aquacultural systems are especially vulnerable, as they are exposed to extreme levels of plastic pollution and provide conditions ideal for disease emergence and spread.

From a One Health perspective, the selection for AMR genes in non-clinical settings may pose a serious risk to human health. Although, the most strongly enriched AMR genes in the present study were generally found in species not known to be human pathogens, there is potential for horizontal transfer events to move these genes into species of clinical relevance. Indeed, environmental bacteria can not only act as vectors for the transmission of AMR genes, but also as their sources. Thus, the addition of selective forces that drive the enrichment of AMR genes in environmental settings can contribute to their biogeographic expansion. Such processes need only point sources of AMR genes to have global consequences.

Given the widespread problem of plastic waste entering the environment, the consequent enrichment of AMR and pathogenic traits is likely occurring in polluted sites worldwide. Such changes pose an interconnected risk to plant, animal, and human health, with the potential to further fuel the global resistance crisis and increase the total burden of disease among marine macroorganisms.

## Acknowledgements

This work was supported by funding from the Australian Research Council to ST (#DE150100009) and IP (#FL140100021).

## Author contributions

ST and EV designed the study. EV, AF, and TG performed the data analyses. All authors contributed to data interpretation, writing the original draft, and reviewed and edited the final draft.

## Notes

### Competing Interest Statement

The authors have declared no competing interest.

