## Supplementary Fig. 1 for "Plastic Leachate Exposure Drives Antibiotic Resistance and Virulence in Marine Bacterial Communities"

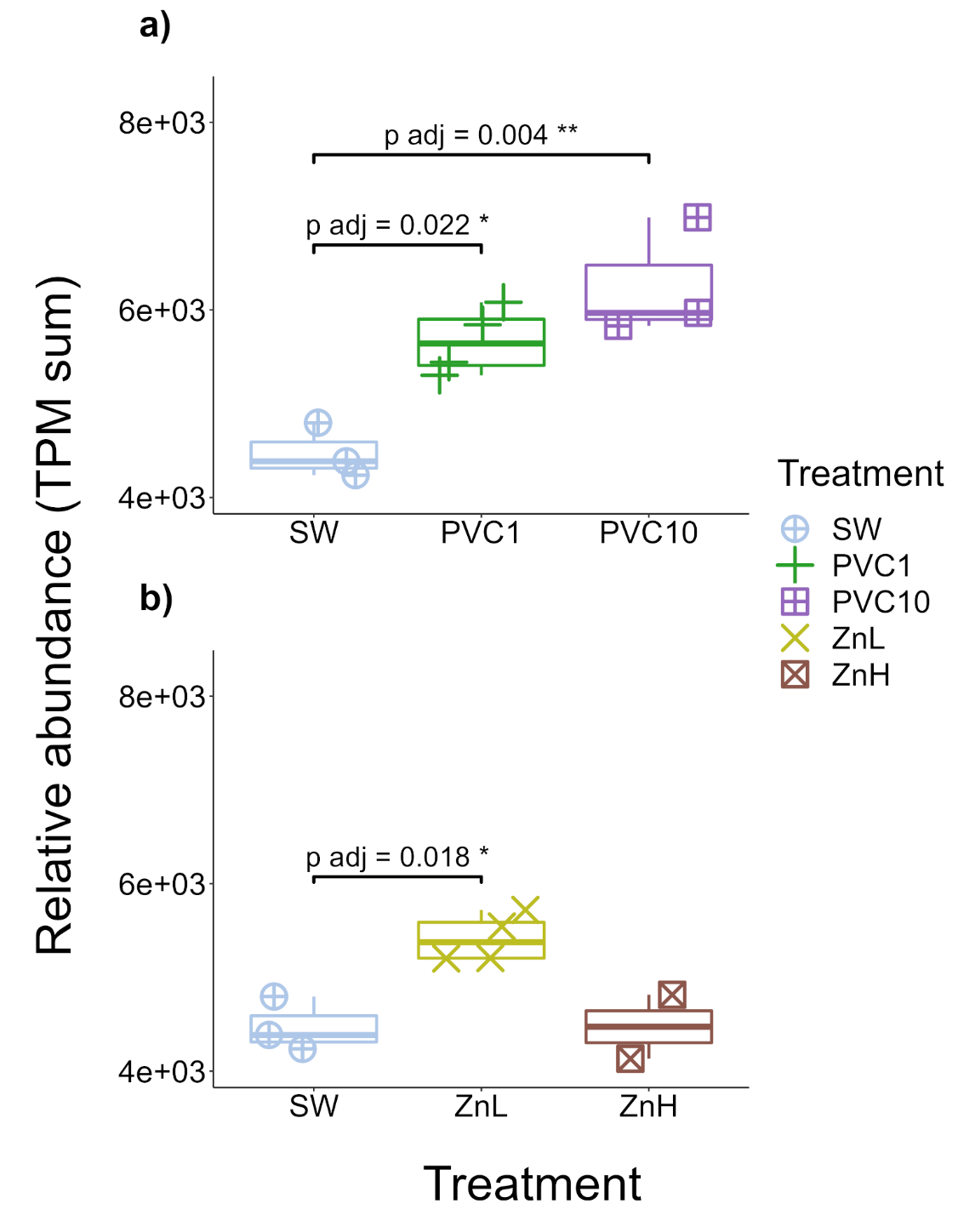

**Suppl. Fig. 1**. Relative abundance (TPM sum) of genes classified as carrying Virulence as predicted by SeqScreen for (a) 1% PVC leachate (PVC1), 10% PVC leachate (PVC10), and (b) 0.13 mg/L Zinc chloride (ZnL), and 1.3 mg/mL Zinc chloride (ZnH) treatments, compared with seawater (SW). Tukey-HSD adjusted p-values have been reported for treatments which differ significantly from the control. The full set of statistical results for all tests are provided in Supplementary Table 5(b-e).
